# Effective sequence-to-expression prediction for a model membrane protein using machine learning and computational protein design

**DOI:** 10.1101/2025.09.25.678317

**Authors:** Yuxin Shen, Maddie Lewis, Juno Underhill, Adrian J Mulholland, Diego A Oyarzún, Paul Curnow

## Abstract

The recombinant expression of integral membrane proteins is notoriously challenging. One way to address this challenge is via computational genotype-to-phenotype models that determine how particular sequence features correlate with protein expression levels. However, the potential of such approaches is yet to be fully realised, at least partly because so few expression datasets are available. Here, we study the sequence-to-expression relationships of a library of 12,248 variants of a specific membrane protein derived from combinatorial computational design. The major advantage of this approach lies in the controlled sequence diversity explored in design, making this new dataset directly compatible with lightweight off-the-shelf bioinformatic tools. The expression phenotype of the entire library is assessed in the widely-used recombinant host *Escherichia coli*. We employed a relatively small dataset of ∼2000 phenotyped sequences to train a sequence-to-expression predictor using supervised machine learning, which achieved high classification accuracy on held-out test sequences. This model was then used to infer the expression of >10,000 unmeasured sequences, and validation of the top predictions of both high and low expressers achieved 100% success rate. Using tools from explainable AI, we identified specific sequence positions and substitutions that are most important in dictating cellular expression levels. This analysis was validated by model-guided protein engineering that achieved an 8-fold increase in the purification yield of a poorly-expressing variant. Our results show that, at least for this controlled dataset, straightforward and interpretable machine learning can reveal the intrinsic sequence code for membrane protein expression.

## Introduction

Integral membrane proteins constitute ∼20% of all proteins and are critically important for cellular life. However, the study of membrane proteins remains plagued by a well-attested and long-standing problem: Why are so few of these proteins tolerated by recombinant hosts? The inadequate (or entirely failed) recombinant production of many membrane proteins remains a major bottleneck for both fundamental research and industrial applications. The interlinked stages of recombinant production - gene expression, membrane localization, bilayer insertion and protein folding - are all potential points of failure, and structural genomics programmes have reported expression failure rates of >90% [1]. Yet the host organism must routinely biosynthesise its own membrane proteins, including some at high abundance, and high-level recombinant expression can certainly be achieved in some cases [1, 2]. This implies that there is specific information encoded within a gene or protein sequence that somehow underpins successful membrane protein production. Structural biology, enzymology, synthetic biology, biomanufacturing, and bioprocessing would all benefit from an improved understanding of the sequence factors that underpin the consistent, reproducible and high-level recombinant production of integral membrane proteins.

In principle, the intrinsic sequence code controlling recombinant membrane protein production could be deciphered by machine learning to produce analytical sequence- to-expression models [3]. Such models would ultimately allow investigators to, for example, pre-select expression-compatible homologues from genomic databases and evaluate the likely impact of specific mutations on protein abundance [4]. However, a universal model remains far beyond reach because sequence-to-expression models have not been found to generalise beyond very specific training conditions [5]. This is almost certain to apply to existing models developed for membrane proteins which use sequence encodings from collected sequence features [6], biophysical-structural and evolutionary properties [7, 8] and codon choice [9]. The major issue lies not in the sophistication of such models but in a deficiency of data; the available training sets for membrane proteins are relatively small and do not necessarily have characteristics that are compatible with machine learning [3]. Since a general model appears unachievable in the immediate future, a more productive path is to develop task-specific models that focus on a particular target protein. However, so far, it is still unclear whether this can be achieved with common, off-the-shelf public ML models or whether bespoke and tailored approaches will be required.

Here, we introduce a strategy for understanding and optimising membrane protein expression in the recombinant workhorse *Escherichia (E.) coli* [10] that exploits large variant libraries generated by computational sequence design. This approach draws upon our own recent description of a first-in-class *de novo* integral membrane protein [11]. We designed a small, heme-binding membrane protein that was inspired by the heme centres that facilitate electron transport in respiration and photosynthesis. To this end we used computational tools to transform a soluble di-heme four-helix bundle [12] into a membrane protein by ‘surface-swapping’; that is, the computational substitution of solvent-facing residues to hydrophobicize the protein exterior. During this sequence design the residues involved in critical protein-protein and protein-heme interactions were strictly maintained, while 54 exterior residues were allowed to sample a restricted hydrophobic amino acid alphabet within an implicit membrane energy function. This surface-swapping approach generated thousands of individual sequence designs (decoys) and a few of these protein sequences were selected for genetic encoding and further characterisation. We found that one of these designed sequences, which we named CytbX, was successfully expressed in recombinant *E. coli* and could be isolated from the cell inner membrane in the hemoprotein form [11]. Other variants from the same design run were similarly well-folded but were expressed at lower levels in the cell.

We reasoned that this novel design library could be used as a testbed for building genotype-phenotype models of membrane protein expression, because the controlled combinatorial diversity between these designs is balanced by tight constraints over variables such as sequence length, codon usage, number of transmembrane (TM) helices and loop composition. The design scaffold captures several characteristics of natural transmembrane proteins including a complex multipass topology and the binding of bioenergetic cofactors, and a major advantage is that the library proteins are short (113 amino acids) and so can be encoded by multiplexed synthetic genes. To provide an expression readout these *de novo* proteins can be genetically fused to green fluorescent protein (GFP), an approach that has been deployed at scale to rapidly phenotype and sort very large protein libraries [13] and is widely used to measure the recombinant expression of diverse sets of integral membrane proteins [2, 14–21]. While *functional* expression can be different from *total* expression [8, 22], high-expressing GFP fusions have been taken as a good starting point for molecular studies [23].

Here we show that, for this novel dataset, established methods for assumption-free and mechanism-agnostic machine learning can discern the sequence basis of membrane protein expression and accurately infer the expression levels of unseen variants. Experimental validation reveals that the expression code learned by the model for this protein is context-dependent and, because of the nature of the training data, is sensitive to the presence of the GFP fusion. This finding in particular may have important implications for future studies seeking to combine multiple expression datasets which use dissimilar expression constructs. Explainability analysis and rational mutagenesis disclose that expression can be determined by just a few key residues. Overall we demonstrate that, given appropriate data, accessible machine learning tools can provide highly accurate sequence-to-expression models for a recombinant membrane protein.

## Methods

### Library design

The 18,339 original design decoys produced in Hardy et al [11] were unduplicated using the seqkit function rmdup. The resulting 12,248 unique protein sequences were backtranslated to DNA sequences with the EMBOSS tool backtranseq using the *E. coli* codon usage table for high-expressing proteins (*Eecoli_high.cut*). Universal adapters were added to the upstream end (5’-ACTTTAAGAAGGAGATATACCATG) and downstream end (GCGGCCGCACTCGAGCTGGTGCCGCGCGGCAGCA-3’) of each sequence to introduce a transcriptional start codon and to facilitate cloning and amplification. All 12,248 gene sequences were ordered as a combined oligo pool from Twist Bioscience.

For diversification of the supplied oligo pool, a mixed-template PCR reaction was performed with the degenerate primers 5’-ACTTTAAGAAGGAGATATACCATG-3’ (Forward) and 5’-GGCACCAGCTCGAGTG-3’ (Reverse) targeting the universal adaptor sequences at either end of all pool genes. Phusion polymerase (New England Biolabs), which has a processivity of ∼100nt, was used in the supplied high-fidelity buffer with 0.2 mM dNTPs, 0.5μM each primer, 3% DMSO and approximately 1ng of the oligo pool. The reaction mixture was transferred from ice to a pre-heated block at 98°C for 30s prior to 10 cycles of 98°C for 10 s, 59°C for 30s, 72°C for 15s, with a final extension at 72°C for 5 min. The 383 bp amplicon was excised from a 2% Tris-acetate-EDTA (TAE)-buffered agarose gel and recovered using the QIAgen gel extraction kit.

### Library construction

Library cloning used restriction/ligation. The diversified oligo pool (12 ng) was digested with 10 units each of NcoI-HF and XhoI (New England Biolabs) in the manufacturer’s SureCut buffer at 37°C for 1h. This digested fragment was recovered using the NEB Monarch PCR purification kit using an elution volume of 10μl. An in-house pET28-GFP plasmid [11, 24] was prepared with NcoI-HF/XhoI using 2μg plasmid and 20U of each enzyme. After 30 min at 37°C, the digested plasmid was dephosphorylated with 5U NEB Antarctic Phosphatase for 30 min at 37°C. The digested plasmid was recovered from a 1.8% agarose TAE gel using a QIAgen gel extraction kit, eluting in 30μl volume. DNA concentrations were determined with a Qubit 4 instrument (ThermoFisher). 40 ng of the digested dephosphorylated vector and 8 ng of the digested oligo pool, constituting 20:60 fmol ends or 1:3 plasmid:insert mole ratio, were ligated overnight on ice with T4 ligase (NEB M0202). The entire ligation mix was dialysed on the shiny side of a 25 mm MCE 0.025 μm filter paper (Whatman VSWP02500) floating in a petri dish of deionised water for 90 mins. The dialysed sample was recovered and 10 μl was transformed into 80 μl 10Beta ultracompetent *Escherichia coli* cells (New England Biolabs) via electroporation using a BioRad MicroPulser instrument with a 0.1cm cuvette on the instrument setting ‘Ec1’ (1.8 kV). After 1h outgrowth in 975 μl of the recovery broth supplied with the competent cells, the library transformation was transferred to 100 ml LB broth plus 50 μg/ml kanamycin (LB/Kan) for selection of transformed cells. A sample of the transformed cells was plated onto LB/Kan agar and colony counting determined ∼80,000 CFUs from the transformation. Subsequent DNA sequencing of selected colonies gave an approximate library size of ∼27,000 unique sequences. After overnight growth of the 100 ml broth culture the library was recovered by plasmid miniprep and transformed into the *E. coli* expression strain BL21(DE3) C43 [25] with at least 500,000 transformants at this stage. The library was stored in 1 ml aliquots in 15% glycerol at −80°C.

### Sanger DNA sequencing

Sanger sequencing of random colonies from the cloned library was performed in a single direction only with the commercial standard sequencing primer *petup* (5’-ATGCGTCCGGCGTAGA-3’; EurofinsGenomics). Reads were of sufficient length to cover the entire library insert and confirm reading frame congruence with the GFP fusion.

### Cytometry analysis and cell sorting

For flow cytometry, 1 ml of frozen library stock was used to inoculate 100 ml of LB/Kan in 250 ml baffled glass flask with a foam bung. Cultures were grown to A600 of 0.8 at 37°C with rotary shaking at 220 rpm before induction for 2 h with 25 mg/l isopropyl-β-D-thiogalactopyranoside (IPTG). The media was also supplemented with 25 mg/l of the heme precursor δ-aminolevulinic acid (ALA) upon induction. A 100 μl sample was removed from the culture and diluted into 10 ml filter-sterile PBS to give ∼10^6^ cells/ml. Cytometry was performed at the Flow Cytometry facility of the University of Bristol using a FACSAria instrument with a 70 μm nozzle. DRAQ7 was used to determine cell viability and side-scatter was recorded with a violet laser (405/20). A negative control (CytbX without GFP) and a positive control (CytbX-GFP as published [11]) were cultured alongside the library and used for comparison.

Cytometry analysis was performed with 500,000 DRAQ7-negative GFP-positive singlet cells (∼6x library coverage). Two gates were defined for the unsorted library to incorporate approximately 8% of the total population at either tail of the distribution. This deliberately excluded a small proportion of very bright clones that were GFP-only cloning artefacts. 10,000 individual events were collected from each gate. The selected cell populations were re-grown and sorting was repeated twice.

The size and diversity of the sorted libraries was determined by colony counting and sequencing. The presumed low-expressing ‘dim’ and higher-expressing ‘bright’ libraries constituted ∼3000 and ∼8000 viable colony-forming units (CFUs) respectively after sorting. For calculating next-generation sequencing (NGS) coverage this was conservatively assumed to be the size of the library *i.e*. 3000 or 8000 individual sequences, although it was expected that most library members would be represented more than once among the CFUs. The sorted libraries were independently sequenced with an Oxford Nanopore MinION device using a rapid sequencing kit (SQK-RAD114) and R10.4.1 flow cell (FLO-MIN114.001). Sequencing over 21h was sufficient for >2 Gbp, giving at last 30x library coverage. Postrun basecalling was performed with Dorado 0.7.1 in duplex mode on the University of Bristol high-performance computing cluster BluePebble1, using the super-accurate model dna_r10.4.1_e8.2_400bps_sup@v5.0.0 with default trimming of sequencing adapters. Sequencing calls for the ‘dim’ and ‘bright’ library featured 11% and 16% duplex sequences, respectively, with N50 at the full plasmid length of 6.4k. As expected, a proportion of plasmid dimers were evident to the same extent in both sorted libraries. After removing the first 150nt of every read, which were found to be of lower average quality, the sequencing data were filtered to select full-length plasmid reads with read quality ≥Q30 using seqkit [26] and fastq-filter (https://github.com/LUMC/fastq-filter). Open reading frames for the library-GFP genes were extracted by EMBOSS getorf [27] and truncated to remove the GFP fusion. This resulted in 9407 library gene sequences for the ‘dim’ library and 7529 sequences for the ‘bright’ library. As expected, at least 70% of these individual gene sequences were duplicated at least once within each library. Deduplication and manual removal of a small number of cloning artefacts provided a final set of 1054 unique low-expressing sequences and 1122 unique high-expressing sequences. For data analysis, FastQC and Unique.seq were accessed via the GalaxyEU webserver (https://usegalaxy.eu) [28]. Only 4 sequences were found to be shared between the two libraries. About one-third of the sequences in each library population came from the original designed oligo pool, meaning that about two-thirds were diversified sequences produced via mixed-template PCR.

### Bulk fluorescence measurements

For bulk expression measurements, individual library members in strain C43 were grown overnight at 37°C/220 rpm in LB/Kan in 10 ml glass flat-bottomed universal vials. 1 ml of each overnight culture was inoculated into 100 ml LB/Kan in a 250 ml baffled glass flask with a foam bung and grown on as above. After reaching an A600 of 0.7 cultures were induced with IPTG with ALA supplementation as above. After 3h post-induction growth, cultures were diluted 17x into PBS and the GFP fluorescence at 512 nm was determined immediately in a Cary Eclipse fluorimeter with excitation at 490 nm.

### Protein purification

In all cases expression was in strain C43. A single colony was picked from a transformant plate into 100 ml LB/Kan in a baffled 250 ml glass flask with a foam bung. This primary culture was grown overnight at 37°C in a shaking incubator at 220 rpm. For secondary expression cultures, 10 ml of the primary culture was used to inoculate each of three volumes of 1 l LB/Kan in 2.5 l baffled glass flasks. This was grown on at 37°C/220 rpm to an A600 ∼0.7, at which point protein expression was induced with 25 mg/l IPTG/ALA as above. After 3h post-induction growth the cells were harvested by centrifugation and purification of individual proteins was performed as described [11]. Briefly, harvested cell pellets were resuspended in total volume 100ml PBS and lysed in a continuous flow cell disruptor at 25 KPSI. Membrane fragments were isolated from the cell lysate at 170,000 x g and solubilised with 1% 5-Cyclohexyl-1-Pentyl-β-D-maltopyranoside (Cymal-5) in 50 mM sodium phosphate buffer pH 7.4, 150 mM NaCl and 5% glycerol. After removing insoluble material via centrifugation at 170,000 x g the detergent-solubilised protein was purified in the same buffer with 0.24% Cymal-5 on a 5 ml Ni-NTA column at a flow rate of 3 ml/min. Sample loading was at 20 mM imidazole, column washing used 4 column volumes with 75 mM imidazole, and elution was performed with 2 column volumes of 0.5 M imidazole at a flow rate of 1 ml/min. The sample was adjusted to 2.5 ml and immediately desalted using a CentriPure Zetadex-25 column under gravity flow. The concentration of the purified sample was determined by absorbance spectroscopy, using an extinction coefficient for the oxidised heme peak at 418 nm of 155,200 M^-1^cm^-1^ [11]. Cellular protein abundance was estimated assuming ∼3x10^6^ proteins per cell [29] and ∼5x10^5^ inner membrane proteins per cell [30].

### In-gel fluorescence

For in-gel fluorescence, induced cultures were harvested and resuspended at 20x in 1:2 PBS:BugBuster reagent plus DNAse. After 45 min at room temperature, SDS-PAGE gel loading buffer (ThermoFisher NP0007) was added to 1x and samples were either used immediately or boiled at 95°C for 5 mins before loading. Sample loading was corrected for the total lysate protein concentration determined using a BCA assay in 2% SDS. Gel imaging used 488 nm excitation on a Typhoon instrument (GE Healthcare).

### RT-qPCR

Methods for RT-qPCR are reported according to the MIQE guidelines [31]. A total RNA cell extract from ∼1x10^9^ *E. coli* cells was isolated from induced and uninduced cultures with the Monarch Total RNA Miniprep kit (New England Biolabs, T2110), incorporating lysozyme for cell lysis and DNase treatment according to the manufacturer’s instructions. The extract was immediately frozen at −80°C and stored for <1 month, only being thawed on ice immediately prior to use. Total RNA was determined with a Qubit 4 instrument (ThermoFisher Scientific; HS RNA, Q32852), also used to confirm <5% DNA in each extract, and 50ng total RNA was used in each reaction. Biological replicates of two duplicates each were performed using the same primer pair (Forward: 5’-ATGGGTTCTCCGTGGCTG-3’, Reverse: 5’-AGACCGGTCAGGAAAACCAG-3’; EurofinsGenomics) generating an amplicon of 239 bp that differs by only two nucleotides between the two variants studied. Data were normalized against *cysG*, *idnT* and *hcaT* as suggested by Zhou et al [32], using the primers from their study, with all reference reactions performed in parallel to the test genes. The reaction was performed with SYBR fluorescent probe detection using a commercial one-step RT-qPCR kit (New England Biolabs Luna Universal kit, E3005) according to the default cycling parameters and aliquoting scheme recommended by the manufacturer. Reactions were 20 μl volume in 96-well plates sealed with adhesive film (MicroAmp N8010560 and 4360954, Applied Biosystems). Thermal cycling was 10 min at 55°C, 1 min at 95°C, 40 cycles of 10 s at 95°C and 30-60 s at 60°C, immediately followed by a melt of 60°C-95°C at 0.15 °C/s. All pipetting was performed manually by Paul Curnow. Data were collected on a QuantStudio3 instrument in fast mode with provided analysis software v. 1.5.3 (Applied Biosystems). Amplification specificity was determined by post-reaction agarose gel, which included no-mRNA and no-transcriptase controls. An uninduced culture of the low-expressing variant was used as the reference for calculating ΔΔCt, with Ct <30 in all samples, a linear dynamic range between 3-1500 ng total RNA, and efficiency ≍1. The baseline and threshold values were calculated by the instrument and used without manual adjustment.

### Machine learning

A random forest model was implemented using scikit-learn (https://scikit-learn.org) for expression classification. The amino acid sequences were assigned binary categorical labels according to their expression level (1=high-expressing, 0=low-expressing), and randomly shuffled prior to training. The amino acid sequences went through one-hot encoding and were split into training (80%) and testing (20%) sets *before the one-hot matrices were flattened into 1D vectors and scaled using StandardScaler prior to 5-fold cross-validation.* The random forest model was trained with n_estimator = 100, min_samples_split = 2 and min_samples_leaf = 1. The trained model was used to infer the expression level of 11,503 unseen protein sequences encoded by the original oligo library. For each sequence, the model outputs the predicted probability for the sequence to be in the bright (high-expressing) class, which was used as an indicator of predicted sequence performance.

Feature importance values were obtained directly from the random forest model based on the mean decrease in impurity during training. For each sequence position, importance scores of different amino acids were aggregated to yield a single positional importance value across the 113 positions. Additionally, Shapley additive explanations (SHAP) analysis was performed on the trained model using 1,000 of the unseen protein sequences to assess the contribution of each amino acid at each position.

### Synthesis and purification of selected library sequences

Sequences with predicted high or low expression characteristics were obtained as individual synthetic genes from Twist Bioscience, retaining the codon usage of the original library and incorporating a His10 tag at the C-terminus. For subcloning to the GFP vector, these genes were amplified by PCR using the universal reverse primer 5’-ATGATGCTCGAGTGCGGCCG-3’ and one of three forward primers as appropriate: 5’-CTTCGACCATGGGTTCTCCGATCCTGCGT-3’, 5’-CTTCGACCATGGGTTCTCCGTGGCTGCGT-3’ or 5’-CTTCGACCATGGGTTCTCCGATCATCCG-3’. The individual amplicons and the GFP destination vector were digested with NcoI-HF/NotI and the vector was dephosphorylated prior to ligation with a rapid ligase (New England Biolabs M2200) for 10 min at room temperature before transforming into chemically-competent TOP10 *E. coli* cells (ThermoFisher). All constructs were confirmed by sequencing.

## Results

An overview of the experimental approach used here is shown in Figure 1. In summary, a library of synthetic oligonucleotides was obtained for 12,248 *de novo* sequences derived from computational protein design, with all of the designed sequences corresponding to a diheme membrane cytochrome comprising four transmembrane α-helices [11]. Codon usage across the oligonucleotide pool was standardised to the usage pattern of high-expressing *E. coli* proteins. These sequences were further diversified by mixed-template PCR and cloned upstream of *gfp* to allow phenotypic selection by fluorescence cytometric cell sorting. Selected clones were sequenced, labelled according to their expression phenotype, and used to train a predictive classifier.

**Figure 1:**
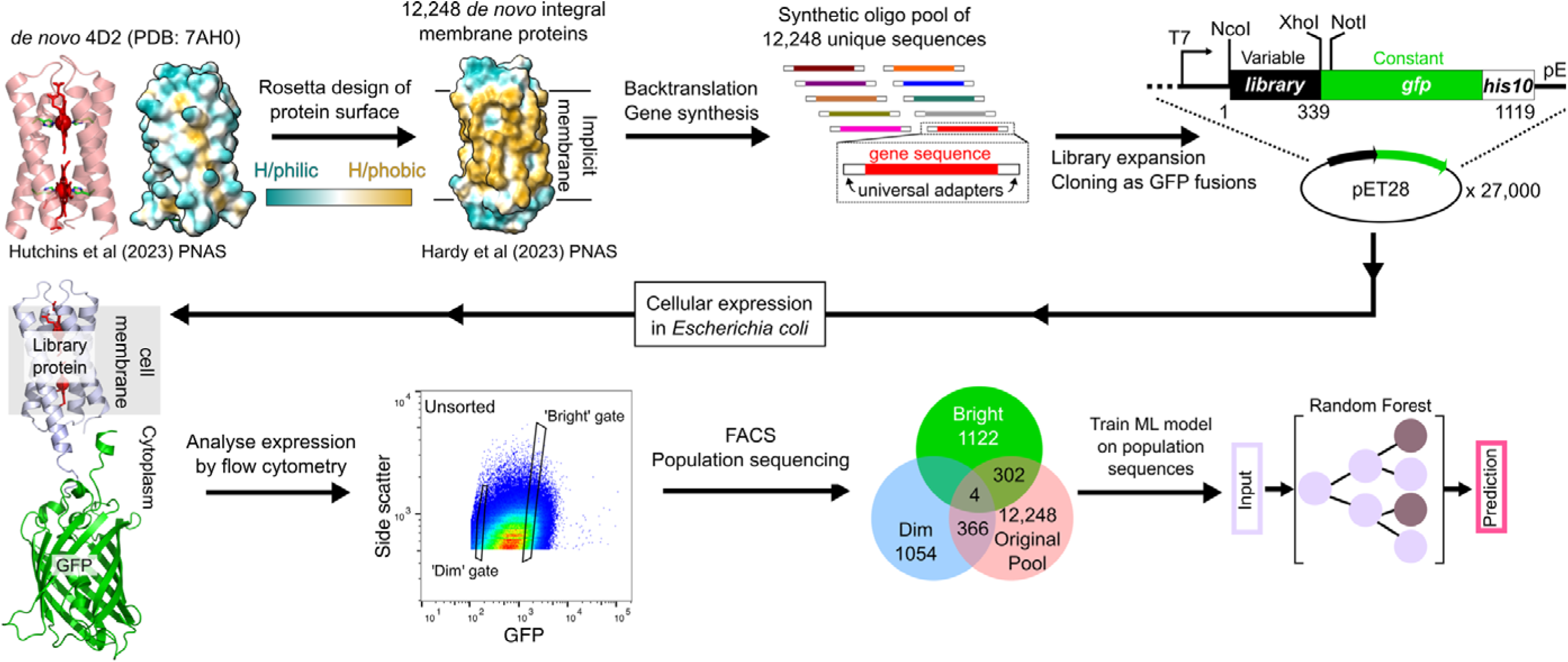
Overview of the experimental approach used in this study. Each of 12,248 computationally-designed membrane proteins were genetically encoded and synthesised commercially as an oligo pool. The combined oligo pool was controllably diversified by PCR and cloned upstream of the open reading frame for GFP. The resulting library was expressed in recombinant bacteria and after flow cytometry analysis, fluorescence-activated cell sorting (FACS) was used to isolate ‘Dim’ and ‘Bright’ cell populations. These two populations were sequenced and the corresponding amino acid sequences used to train a predictive classifier.

### Characteristics of the variant library

The variant library used here is based on a *de novo* protein of our own design. This designed sequence is a multipass integral membrane protein comprising 113 amino acids that coordinates two molecules of heme at the centre of a four-helix bundle, as described in full in ref. 12 and shown in Figure 2. These designer proteins are capable of electron transfer reactions with small molecules and natural enzymes [12] and can be readily purified from *E. coli* membranes in detergent micelles or polymer nanodiscs [33]. The relative simplicity, robustness and sequence versatility [34] of these designs makes them excellent targets for understanding sequence-expression relationships. The original design process stipulated multiple simultaneous amino acid substitutions at 54 sites distributed over the protein surface. Residues at 13 of these sequence positions were allowed to sample a small set of alternative residues while the remaining 41 sites sampled from a minimal amino acid alphabet of FAILVWGST; the Rosetta resfile from ref. 12 specifying the positional alphabet is included here as supplementary data. Evaluation of the resulting 12,248 designed protein sequences revealed that amino acid substitutions occurred throughout the protein (Fig. 2), with some positions showing greater relative diversity (although with low absolute diversity). Analysing a subset of the designed sequences showed a typical between-sequence distance of about 12 residues (Supplementary Figure S1).

**Figure 2:**
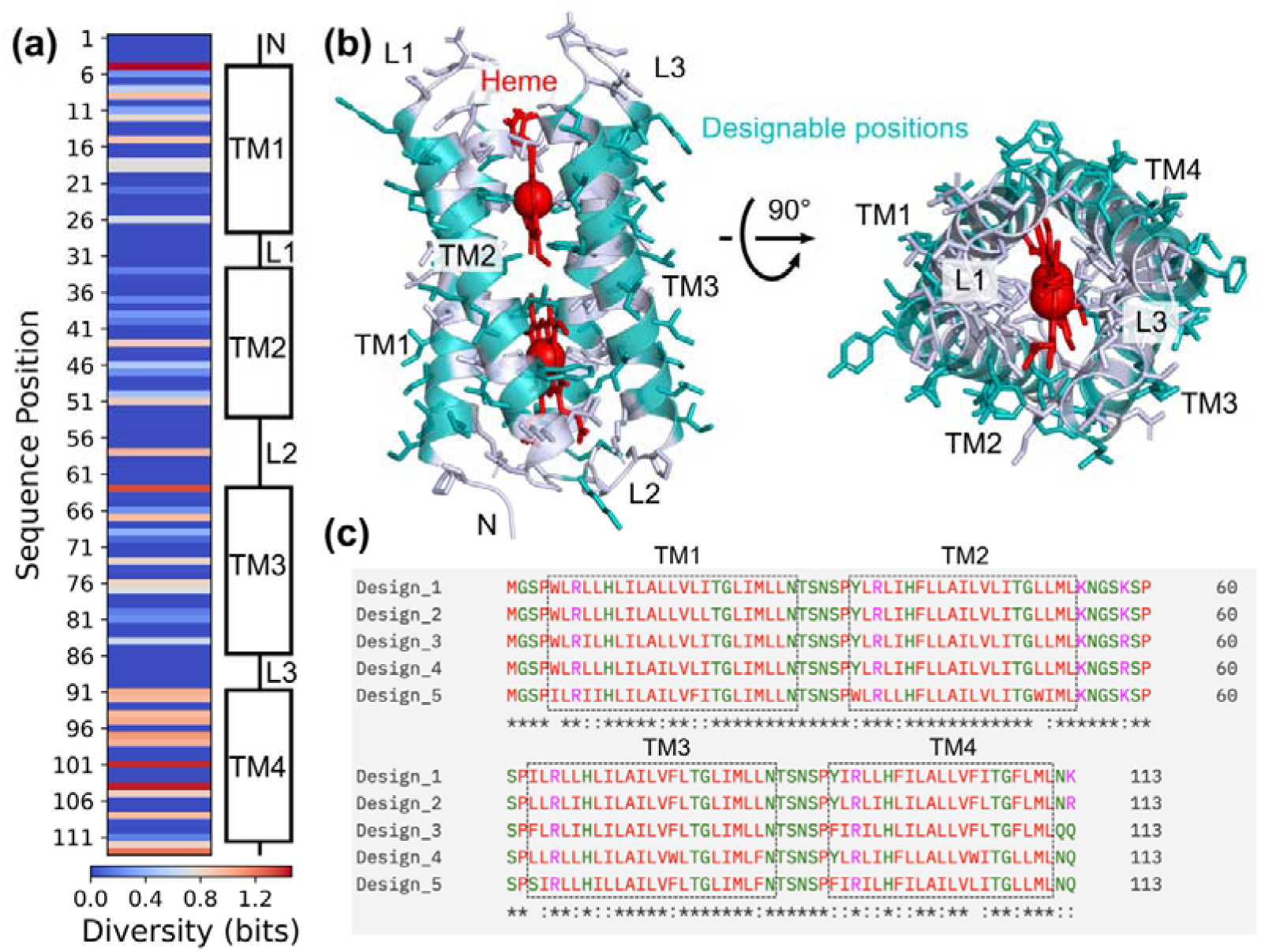
General characteristics of the designed protein library. (**a**) Computationally-designed amino acid substitutions are distributed throughout the diheme four-helix bundle. Some positions showing higher relative diversity, calculated as Shannon entropy. *N*, amino terminus; *TM*, transmembrane; *L*, loop. (**b**) Protein model showing the surface location of the designable positions. (**c**) Illustrative multiple sequence alignment of five sequences highlighting the controlled diversity of the library. Boxed regions correspond to TM helices 1-4 as shown.

### Library preparation

The library variants were obtained as a commercial oligo pool with all oligos using a codon bias associated with high-expressing *E. coli* proteins. This pool was then controllably diversified (shuffled) via mixed-template PCR with a low-processivity polymerase [35] before cloning into a bacterial expression plasmid upstream of the open reading frame for superfolder GFP. This resulted in a fusion protein library with ∼27,000 individual variants. Sequencing a random sample of the library suggested that about 1/3 of the library sequences were from the original oligo pool and the remainder were shuffled versions of the original sequences.

### Phenotypic analysis and sorting

The entire library was transformed into *E. coli* strain BL21(DE3) C43 and cultured for membrane protein expression. Induced cultures were analysed by flow cytometry and sorted into two phenotypic groups using GFP florescence as a selectable phenotype. Our selection strategy was geared towards identifying two distinct sequence populations with ‘high’ and ‘low’ expression characteristics. This was inspired by work showing that simplifying cell sorting data to binary outcomes can give equivalent performance to more complex analyses, including for the prediction of continuous variables such as protein expression [36]. We were not concerned with expression tuning for intermediate levels (which is probably better accomplished by adjusting other parameters such as the strength of ribosome binding sites) but only in identifying intrinsic sequence features that contribute to successful and unsuccessful expression profiles. A more practical motivation was that selecting toward binary phenotypic labels provides efficient data for model training while minimising the costs of sorting and sequencing. Developing lean, cost-effective machine learning methods is likely to be of broadest interest.

After three successive rounds of selection, two populations of dim (low expression) and bright (high expression) phenotypes were isolated from the original library with limited fluorescence overlap between these two library subsets (Fig. 3a). Nanopore sequencing recovered 1122 unique gene sequences (encoding 1066 unique protein sequences) from the sorted bright library and 1054 unique gene sequences (989 unique proteins) from the sorted dim library. Only four gene sequences were found in common between the bright and dim sequence libraries, confirming high enrichment of the respective phenotypic populations. For each of the sorted populations the within-population sequence diversity was comparable to that of the unselected original oligo pool, meaning that the gating strategy was not simply collecting highly similar sequences (Supplementary Figure S1).

**Figure 3.**
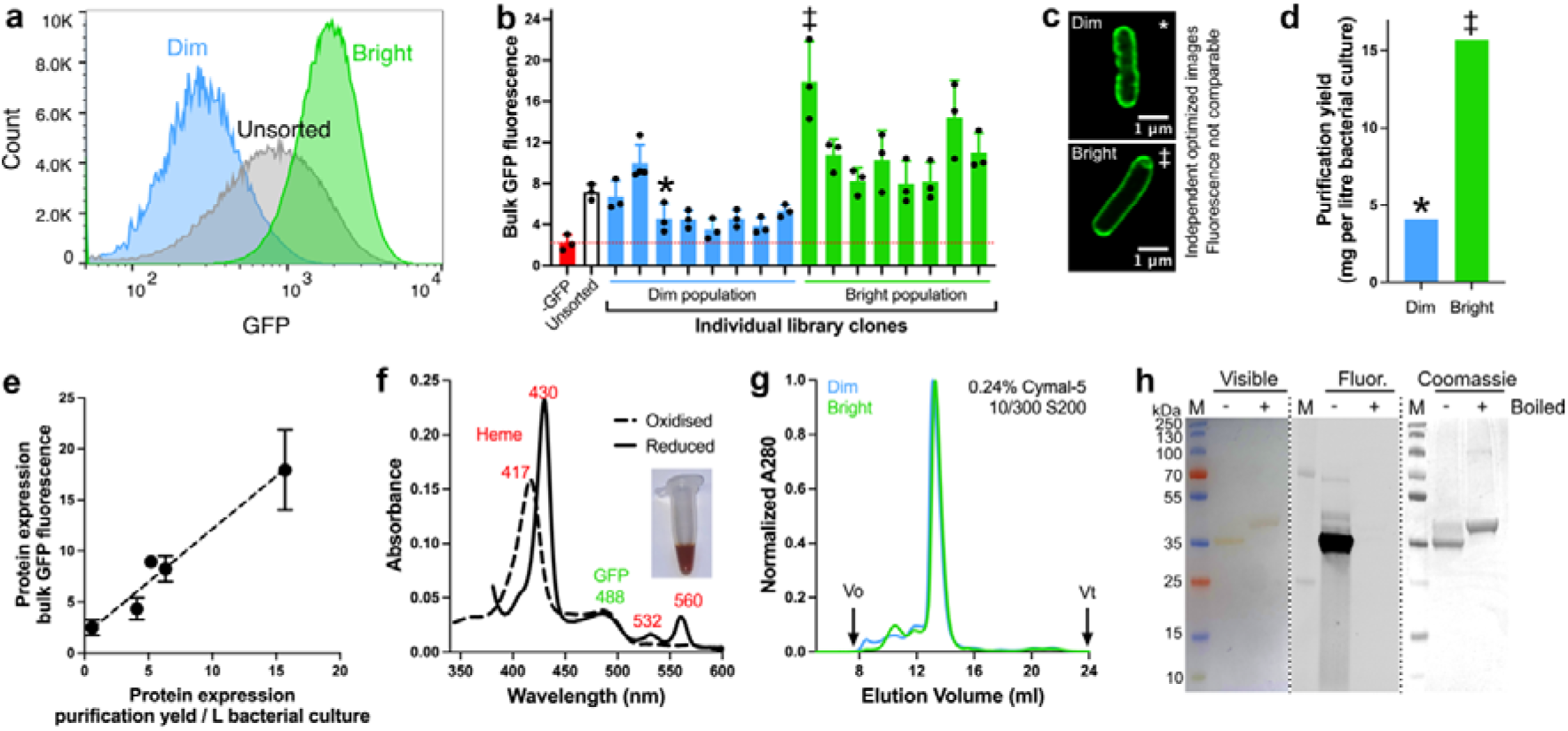
Sorting library-GFP fusion proteins by GFP phenotype and experimental validation of selected proteins. (**a**) Two libraries enriched in low-GFP (dim) or high-GFP (bright) expression phenotypes are selected by three rounds of fluorescence-activated cell sorting. (**b**) Bulk expression measurements of individual selected clones confirm the phenotype associated with each library population. (‡) and (*) denote the same protein across panels b-d. (**c**) Confocal fluorescent microscopy of *E. coli* cells expressing two representative sequences from the dim and bright libraries show that these proteins are localised to the cell membrane. Image contrast is optimised for display and is not representative of the true fluorescence intensity *i.e.* cells expressing the bright protein showed higher GFP under imaging. (**d**) Selected dim and bright individuals are purified by affinity chromatography with different protein yields. (**e**) Purification yields of individual library-GFP proteins correlate directly with their bulk GFP signal. (**f**) Absorbance spectroscopy of purified individual fusion proteins confirms the expected heme cofactor loading and GFP chromophore absorption. Inset photo shows the intense red colouration of the purified protein. (**g**) Size-exclusion chromatography shows that purified dim and bright fusion proteins are monodisperse and have the same apparent size. (**h**) SDS-PAGE analysis confirms the expression of full-length fusion protein. The purified fusion protein remains folded under ambient conditions and a visible coloured band is observed without staining (*Visible*). Sample boiling unfolds GFP, evidenced by a band shift and loss of in-gel fluorescence (*Fluor.*). The heme signal is retained in the visible gel since the designer proteins are thermostable. Coomassie staining (*Coomassie*) shows the high purity of the sample. Theoretical molecular weight of the fusion protein is 42 kDa.

To validate the sorted libraries, a random subset of 8 individual clones from each library population were tested in bulk fluorescence assays where the GFP signal from induced cultures was measured in a fluorimeter. Single clones from the bright library did generally show higher GFP signal than those from the dim library; the dynamic range of these measurements was less pronounced than in cytometry, presumably because of instrument differences and inner filter effects (Fig. 3b). Cell imaging confirmed that the fusion proteins were localized to the cell membrane, and that this membrane-localized protein was the phenotypic readout (Fig. 3c). Individual clones were also purified from cell membrane fractions to confirm that the fluorescence signal correlated directly to the yield of purifiable protein (Fig. 3d and e). This purification used the nonionic surfactant Cymal-5, a mild detergent that does not disaggregate misfolded protein. The yield of the selected library-GFP fusion protein purified from the bright library was an exceptional 16 mg protein per litre of bacterial culture – equivalent to ∼70,000 copies per cell, ∼2% of all cell protein or ∼10% of all inner membrane protein – confirming that the library harbours ultra-high-fitness sequences. Absorbance spectroscopy of purified samples determined that the library proteins were fully loaded with heme (Fig. 3f) and size-exclusion chromatography showed that proteins were not disposed to aggregation (Fig. 3g), indicating that misfolding and aggregation propensity *per se* could not explain the difference in expression levels. Finally, gel electrophoresis confirmed the purity and integrity of the isolated protein (Fig. 3h and Supplementary Figure S2).

### Predictive sequence-to-expression model

To assess the suitability of the selected libraries for machine learning we assigned categorical labels of either 1 (bright; high expression) or 0 (dim; low expression) and used one-hot, ESM2 and ProtBert amino acid sequence representations to train each of four different classifiers: random forest, logistic regression, multilayer perceptron and a transformer. In all cases this used a stratified 80:20 data split for training and testing respectively. Model evaluation on the held-out test set showed that one-hot encoding gave the highest classification accuracy of >0.9 across all models with area under the Receiver Operating Characteristic curve (AUROC) being ≥0.95 (Fig 4a; Supplementary Figure S3). The combination of one-hot encoding with a random forest model was chosen for further work since this offers equivalent performance to the other architectures tested and is compatible with explainability analysis.

**Figure 4:**
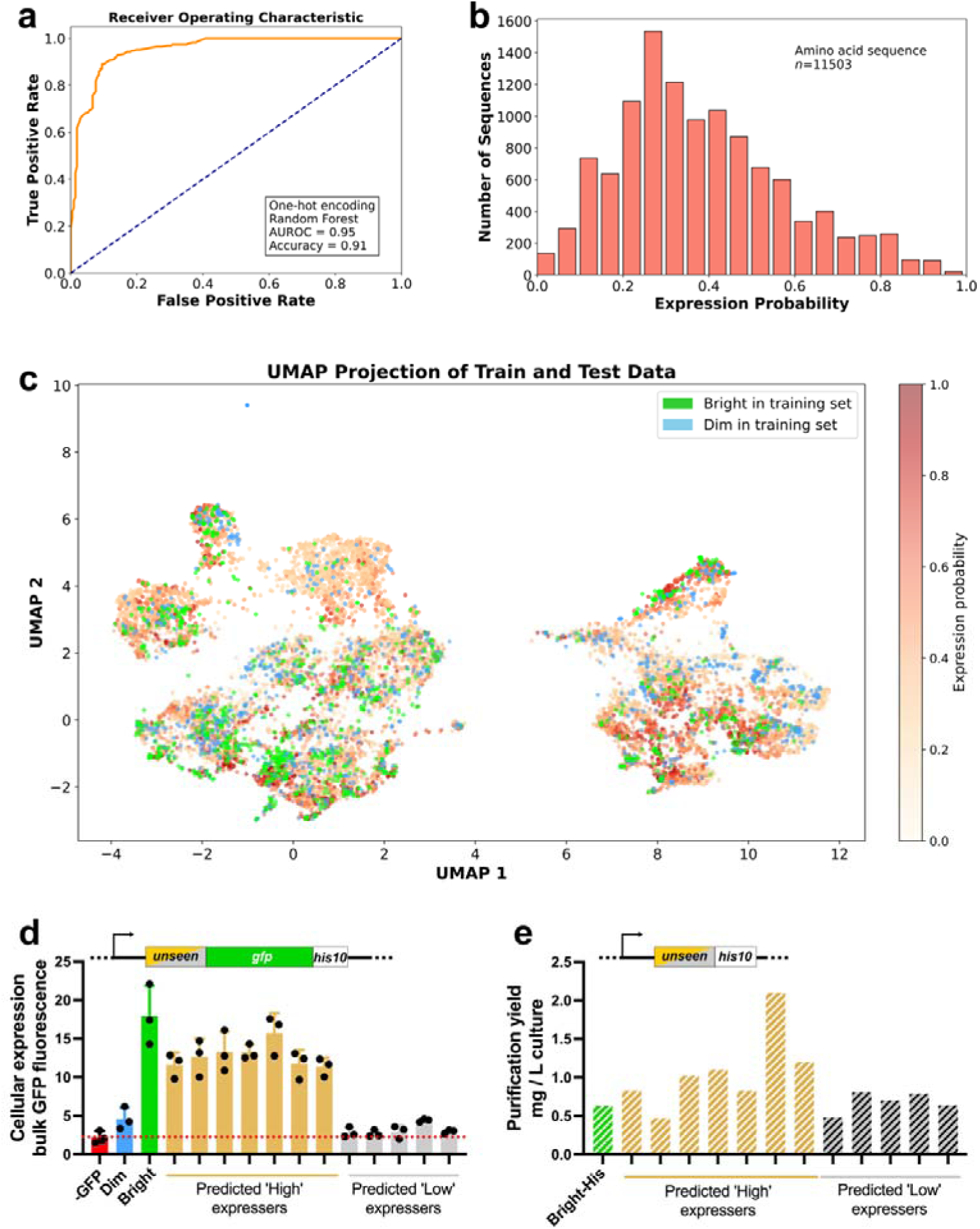
A machine learning model for membrane protein expression. (**a**) Sequence libraries corresponding to either the bright or dim phenotype are labelled discretely, one-hot encoded and used to train a random forest classifier. Testing with 20% of the dataset excluded from training reveals excellent classifier performance. (**b**) The trained model was used to predict the expression phenotype of 11,503 unseen and unlabelled protein sequences encoded by the original oligo pool. (**c**) Dimensionality reduction by UMAP shows two major clusters in the data but no distinct clustering of high- and low-expressing sequences. (**d**) Bulk fluorescence measurements provide experimental confirmation of the phenotypic predictions made by the model. Control samples are from Fig. 1b-d. (**e**) The expression yield and expression profile are sensitive to the C-terminal domain.

The trained random forest model was then queried with 11,503 one-hot encoded unlabelled sequences encoded by the original oligo pool, in order to predict the probability of each variant having the bright phenotype (Fig. 4b). A UMAP representation of both the training set and the unlabelled sequences (Fig. 4c) suggests the existence of two broad and diffuse sequence clusters but with expression levels being distributed across the entire sequence space. This confirms that simple sequence clustering alone cannot be used to explain the expression phenotype. The model predictions also did not correlate with simple biophysical metrics such as the overall Rosetta score, sequence hydrophobicity, or the insertion ΔG of TM1 (Supplementary Fig S4). Sequence representation by one-hot encoding thus appears to capture expression-relevant features that are not easily identified by more intuitive knowledge-based approaches.

For experimental validation of the model predictions we selected a set of 12 unstudied variants with predicted expression scores of <0.02 (high probability of dim) or >0.9 (high probability of bright) and obtained the corresponding genes as individual clones. The observed expression level of these proteins was consistent with the classifier prediction in all cases (Fig. 4d).

Expression datasets for natural membrane proteins have been collected both with [2, 19, 37] and without [1] a GFP tag. Recent studies have now highlighted the importance of the soluble C-terminal domain that follows the final transmembrane helix – the ‘C-tail’ – in controlling membrane protein expression [38, 39]. We were thus motivated to understand whether different C-tail sequences could affect the expression profile of our variants. To determine the sensitivity of these variants to the C-tail, we replaced GFP with a short His10 affinity tag and purified each of these His-tagged proteins by affinity chromatography (Fig 4e). The His-tagged proteins were expressed at significantly lower levels overall and without the clear expression diversity of the equivalent GFP fusions. This result suggests that the ultra-high expression observed in the training data may be restricted to full-length fusion proteins, and that models trained on particular protein constructs cannot necessarily be extrapolated to other contexts. This affirms the strong influence that soluble domains can exert on membrane protein biosynthesis and is an important consideration for further work in this area.

### Expression engineering via explainability analysis

We next used feature importance analysis to identify residues that strongly contribute to expression. Sequence positions 12 and 113 were found to prominently affect the model predictions (Fig 5a). Analysis of these positions with the Shapley additive explanations (SHAP) algorithm for model explainability revealed that position 12 was either F/I/L with I12 being strongly associated with high-expression prediction by the model (Fig 5b). At position 113, library sequences were almost exclusively K or Q and almost all high-expressing sequences featured K113 (Fig. 5b). Much weaker preferences were observed in high-expressing sequences for I/L6, I/F11, I/L19, W/F76, Y91, and I98; low-expressing sequences were marginally enriched in F92, L95 and N/R112.

**Figure 5:**
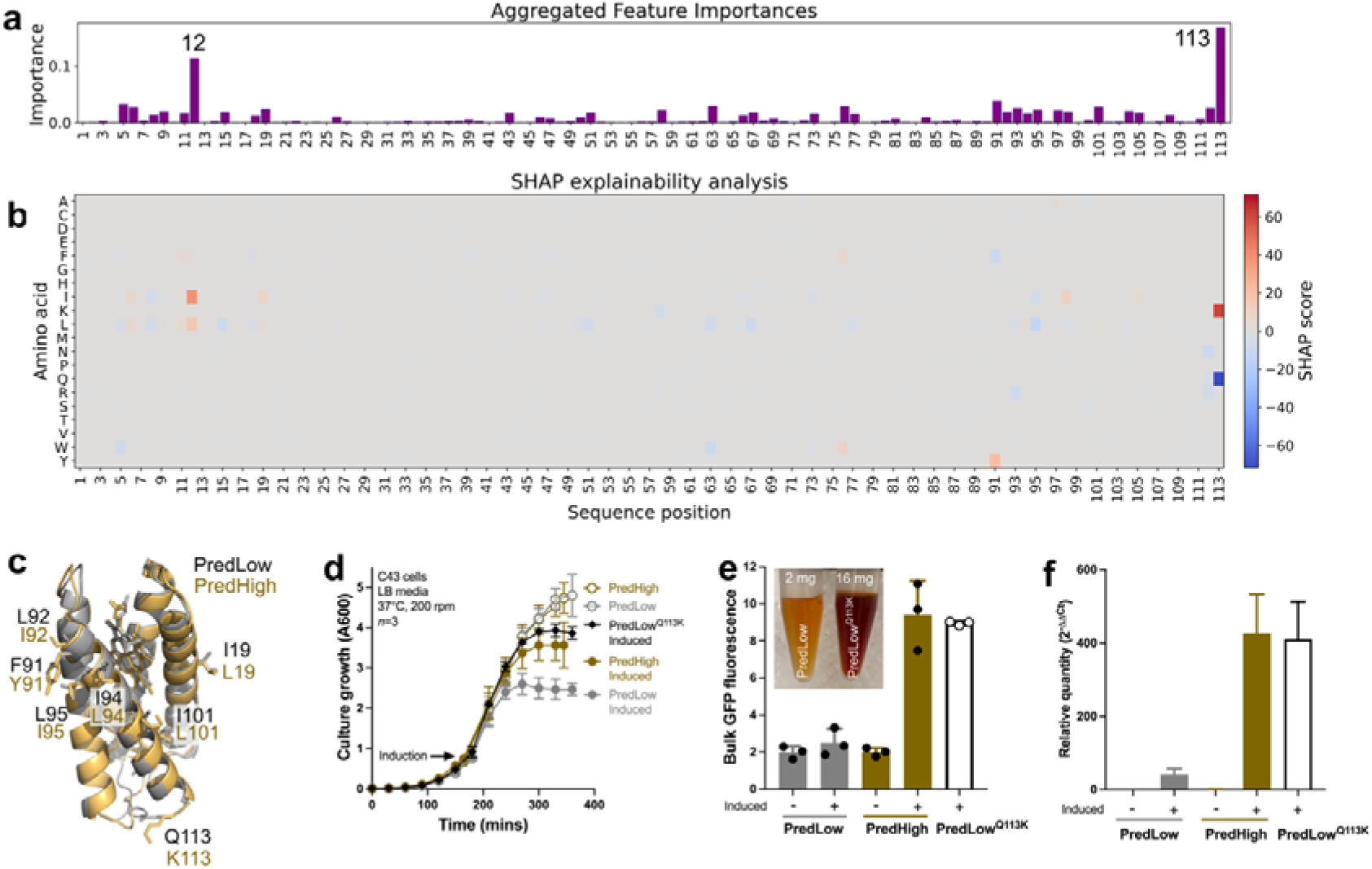
Variant optimization with explainability analysis. (**a**) The classifier identifies two sequence positions, 12 and 113, as being the most important features for determining expression. (**b**) Residue-level SHAP analysis shows that Ile at position 12 and Lys at position 113 are associated with high-expressing sequences; in contrast, Gln 113 is strongly associated with sequences that have low predicted expression. (**c**) Overlay of the AlphaFold structures of two variants that differ by only the 7 amino acids shown but have strongly High or Low predicted expression outcomes, termed *PredHigh* and *PredLow* respectively. (**d**) Growth curves of *E. coli* strains producing PredHigh or PredLow. Cells carrying PredLow show a more pronounced growth arrest after induction. This metabolic burden is completely relieved by a single mutation Q113K. (**e**) Bulk fluorescence measurements demonstrate expression recovery in the engineered variant PredLow^Q113K^. Inset, photograph of purified hemoprotein confirms the associated difference in purification yields as shown. (**f**) Complementary expression analysis by RT-qPCR shows that relative transcript abundance mirrors the bulk fluorescence signal. For additional controls and efficiency curve see Fig. S8.

We reasoned that the predominance of K113 in high-expressing sequences might reflect a specific interaction of the Lys sidechain with the GFP fusion, perhaps allowing the GFP to act as a folding chaperone. To probe this further we purified one of the high-expressing variants in the presence and absence of GFP and directly compared the size-exclusion profile of these two constructs (Supplementary Fig. S5). While the expression yields were dramatically different - being about 30x higher in the presence of GFP – there was no general increase in the aggregation propensity or difference in heme binding of the His-tagged protein. Thus while the presence of the GFP fusion might support folding in the cell, it is not required to maintain the fold of purified library proteins. This conclusion was supported by molecular dynamics simulations, which found that GFP was highly mobile relative to CytbX due to the flexibility of the connecting linker. These simulations could not identify substantive interactions between the two fused proteins but did find that salt bridges sometimes occurred between Lys113 and lipid headgroups (Supplementary Figure S6).

We directly tested the feature importance scores from Fig. 5a-b by investigating two variants with opposite expression outcomes but with very similar sequences, differing by only 7/113 amino acids and 13/339 nucleotides (Fig.5c; Supplementary Fig S7). Both sequences featured Ile at position 12 but were different for K/Q at position 113. Culture growth curves showed that recombinant strains carrying the low-expressing variant (termed PredLow) exhibited a more rapid and pronounced growth arrest after induction versus the high-expressing variant PredHigh (Fig. 5d).

We hypothesised that if position 113 was a critical determinant of expression, then we should be able to substantially improve the expression of PredLow by introducing mutation Q113K (CAG→AAA). This approach was successful, with the rationally engineered variant PredLow^Q113K^ showing minimal growth arrest (Fig. 5d), high bulk fluorescence (Fig. 5e) and an 8-fold increase in purification yield (Fig. 5e). To corroborate these findings we turned to RT-qPCR as an alternative measure for gene expression that is independent of the GFP signal. The relative transcript abundance of PredLow, PredHigh and PredLow^Q113K^ was in direct agreement with the observed fluorescence (Fig. 5f and Supplementary Figure S8).

## Discussion

The results presented here show that the fitness landscape of an integral membrane protein can be successfully described by tractable and explainable machine learning tools. We exploit a novel protein library where multiple simultaneous mutations are introduced computationally at designated surface sites throughout a small *de novo* protein, using a restricted amino acid alphabet and biased by a membrane-specific energy function (Fig. 2). The resulting multipass membrane cytochromes can be fused to a phenotypic marker, are localized to the *E. coli* inner membrane, are straightforward to purify and characterise, and retain a strong heme absorbance signal that reports directly on successful protein folding in the membrane (Fig. 3). The computational search of a structured sequence space provides an alternative to library generation by systematic mutational scanning and random mutagenesis and to the greater diversity of natural protein collections.

Our results confirm that such computationally-generated protein libraries are ideal for machine learning and offer a suitable trade-off between feature diversity and library size (Fig 4). The sequence data are directly compatible with standard off-the-shelf bioinformatics packages and the analysis is of low computational expense, with all classifier training, prediction and feature analysis running in less than 3 minutes on a standard laptop computer. Assumption-free and mechanism-agnostic sequence representations reveal non-obvious features that exert strong control over membrane protein expression but are not necessarily captured by biophysical encoding schemes (Fig. 5). While encodings such as the one-hot representation used here can provide excellent model performance on the specific dataset being studied – often surpassing more intuitive biophysical representations of protein sequence [40, 41] – they do not generalize well [42]. Further work will explore whether hybrid models incorporating mechanistic features will allow extrapolation from our design library to other datasets.

We show that membrane protein expression can be strongly influenced by fusion tags (Fig. 4d-e) and can vary at least 8-fold based on just one or two key residues (Fig 5). In particular, the ultra-high expression of library variants depends upon the presence of Lys at position 113, sited at the end of the last transmembrane helix. We speculate that K113 may be important for specifying protein folding in the cell, and perhaps dictates efficient co-translational membrane insertion and topological definition according to the ‘positive-inside rule’ [43–46]. This idea could be tested in the future by force-unfolding studies to compare the nature of the folding landscape between different variants [47]. We also find that an earlier codon corresponding to Ile at position 12 in TM helix 1 is favoured in, but not exclusively limited to, high-expressing sequences (Fig. 5a). Codon usage, nucleotide composition, amino acid identity and mRNA structure within the early part of the transcript can all be important for prokaryotic gene expression [13, 48, 49], including for membrane proteins [20, 50]. However, whatever positive impact I12 may have on protein expression is evidently secondary to the effect of K113 (Fig. 5). The bias towards I12 in high-expressing sequences remains to be fully understood.

How could the C-terminal region after the last transmembrane helix – the C-tail – so dramatically impact the expression yield (Fig. 4)? Several studies have now identified the importance of the C-tail in controlling membrane protein expression [38, 39, 51, 52] and we have previously found that the production of these *de novo* membrane proteins is enhanced by an extended soluble C-tail [33]. Although a complete analysis lies outside the scope of the current manuscript, we suspect that the length of the tail sequence could be an important factor. Assuming that protein synthesis proceeds via the classical translocon pathway [45, 53], in the early stages of biogenesis the growing nascent chain is transferred to the translocon immediately as it exits the ribosome. Hydrophobic transmembrane sections then move from the translocon into the membrane bilayer, where the final stages of folding and assembly can occur. The distance from the ribosome peptidyl transferase centre to the mouth of the exit tunnel equates to a peptide chain length of about 45 amino acids. A long soluble C-tail - in this case, GFP – means that the ribosome is still engaged with the translocon as the final transmembrane helix leaves the exit tunnel, enabling translocon-mediated co-translational bilayer insertion. In contrast, for a shorter C-tail – such as in the His-tagged constructs used here – peptide synthesis will be complete, and the ribosome dissociated, before the final TM helix has travelled to the translocon. This implies the post-translational insertion of short C-tail proteins [39]. Whatever the precise mechanism may be, our results imply that care should be taken in amalgamating membrane protein expression datasets that have used different tag sequences.

Overall, we have established a new membrane protein expression dataset that will serve as a useful testbed for the development of specific and general sequence-to-expression algorithms. The expression phenotype of these proteins can be interpreted using classical pathways of membrane protein biogenesis. While this study targets a short model protein, we anticipate that the rapid progression of commercial DNA synthesis will soon enable the extension of our method to longer protein sequences. Extending the computational mutagenesis method developed here to a range of other membrane proteins, and to residues within the protein core, should enable the mapping of both local and global sequence parameters that control recombinant expression. Our results also raise the prospect of applying computational resurfacing to natural membrane proteins to achieve enhanced recombinant expression while retaining native structure and function.

## Data availability

The sequence data and code used in this study is available online at https://github.com/Curnow-Lab-University-of-Bristol/Membrane-protein-library-ML. The raw sequencing data is available to the community at the BioStudies repository with accession number S-BSST2184 (https://doi.org/10.6019/S-BSST2184).

## Conflict-of-interest statement

The oligo pool used here was obtained from Twist Bioscience at a heavily reduced price as part of an academic pre-market trial for an enhanced synthesis method. The authors have no affiliation with, or financial interest in, Twist Bioscience. Twist Bioscience was not involved in any other aspect of the research described here or in the preparation of this manuscript.

## Acknowledgements

PC & AJM were supported by the UK Biotechnology and Biological Sciences Research Council (BBSRC) grant number BB/W003449/1. YS was supported by BBSRC grant number BB/T00875X/1. JU was supported by Engineering and Physical Sciences Research Council Doctoral Training Partnership EP/W524414/1. This research used the Flow Cytometry Facility and the computational facilities of the Advanced Computing Research Centre of the University of Bristol (http://www.bristol.ac.uk/acrc/). Thanks to Ben Hardy and Ross Anderson for helpful discussions.

## Author contributions

Conceptualization: PC, DAO. Funding acquisition: PC, DAO, AJM. Investigation: YS, JU, PC. Methodology: YS, JU. Resources: PC, DAO, AJM. Supervision: PC, DAO, AJM. Visualization: YS, JU, PC. Writing – original draft: PC. Writing – review and editing: YS, AJM, DAO, PC. All authors have read and approved the final manuscript.

## Supplementary Information

### SUPPLEMENTARY METHODS

Supplementary methods are provided for model benchmarking in machine learning (Supplementary Figure S3) and the molecular dynamics simulations comprising Supplementary Figure S6.

#### Comparison of feature representations and models

Amino acid sequences were assigned binary categorical labels according to their expression level (1=high-expressing, 0=low-expressing), randomly shuffled and split into 5-fold cross validation. Three classifiers were implemented using scikit-learn (https://scikit-learn.org) for expression classification: a random forest, a logistic regression, and a multilayer perceptron (MLP). The random forest classifier was trained with number of estimators = 100, minimal sample split = 2 and minimum sample leaf = 1. The logistic regression model used L2 regularization with regularization parameter C = 1. The MLP classifier was trained using 1 hidden layer of 100 neurons with ReLU activation. Another Transformer classifier, implemented in PyTorch, consists of a linear embedding layer that projects the input space into 64 neurons in the hidden layer, followed by a single transformer encoder layer with 4 attention heads, and finally a linear layer with sigmoid activation for classification. Three embedding methods were used for each sequence, one-hot (2260 dimensions), ProtBERT (1024 dimensions) and ESM2 (320 dimensions). Each combination of three embeddings and four models were tested with 5-fold cross validation, with the AUROC, accuracy and F1 score shown in Figure X, and all the error bars represent the standard deviation across 5-fold.

#### Molecular Dynamics simulations

MD simulations were performed on the University of Bristol high-performance computing cluster BluePebble. The forcefield was AMBER ffsb14 with TIP3P used for water [1, 2]. Parameters for oxidised heme were derived using the AMBER mdgx procedure. Protein models were predicted using AlphaFold3 [3] and input into a 3:1 DOPG:DOPE lipid bilayer to represent the *E. coli* membrane using PACKMOL-Memgen [4]. These models were solvated in a cubic box with the overall charge of the system balanced by adding the exact number of potassium ions needed to neutralize the system; this was 97 for the low-expressing variant ‘PredLow’ and 98 for the high-expressing variant ‘PredHigh’ (see main text, Fig. 5).

The respective systems were energy-minimized for 1000 steps using steepest descent. This was followed by 1000 steps restraining the protein Cα atoms and a further 1000 steps with no restraints. 100 ps MD was then performed to equilibrate the system with all heavy atoms restrained followed by 100 ps with only Cα restrained. These MD runs were done on GPUs in the NVT ensemble (298K), using SHAKE to constrain bonds containing hydrogen atoms with a timestep of 2 fs for the integration of the equations of motion [5–7]. Long-range electrostatic interactions were calculated using the particle mesh Ewald method with a cut-off of 10 Å for direct contributions. Production simulations were performed in the NPT ensemble (298K, 1 bar) using the Langevin thermostat and the Berendsen barostat [8].

Three replicate 300 ns MD runs were performed for each of the two variants studied. The replicates were initiated with different sets of random velocities. The analysis was performed using the MDanalysis and CPPTRAJ software toolkits and the trajectories were visualized using VMD [9–12].

---

**Supplementary Figure S1.**
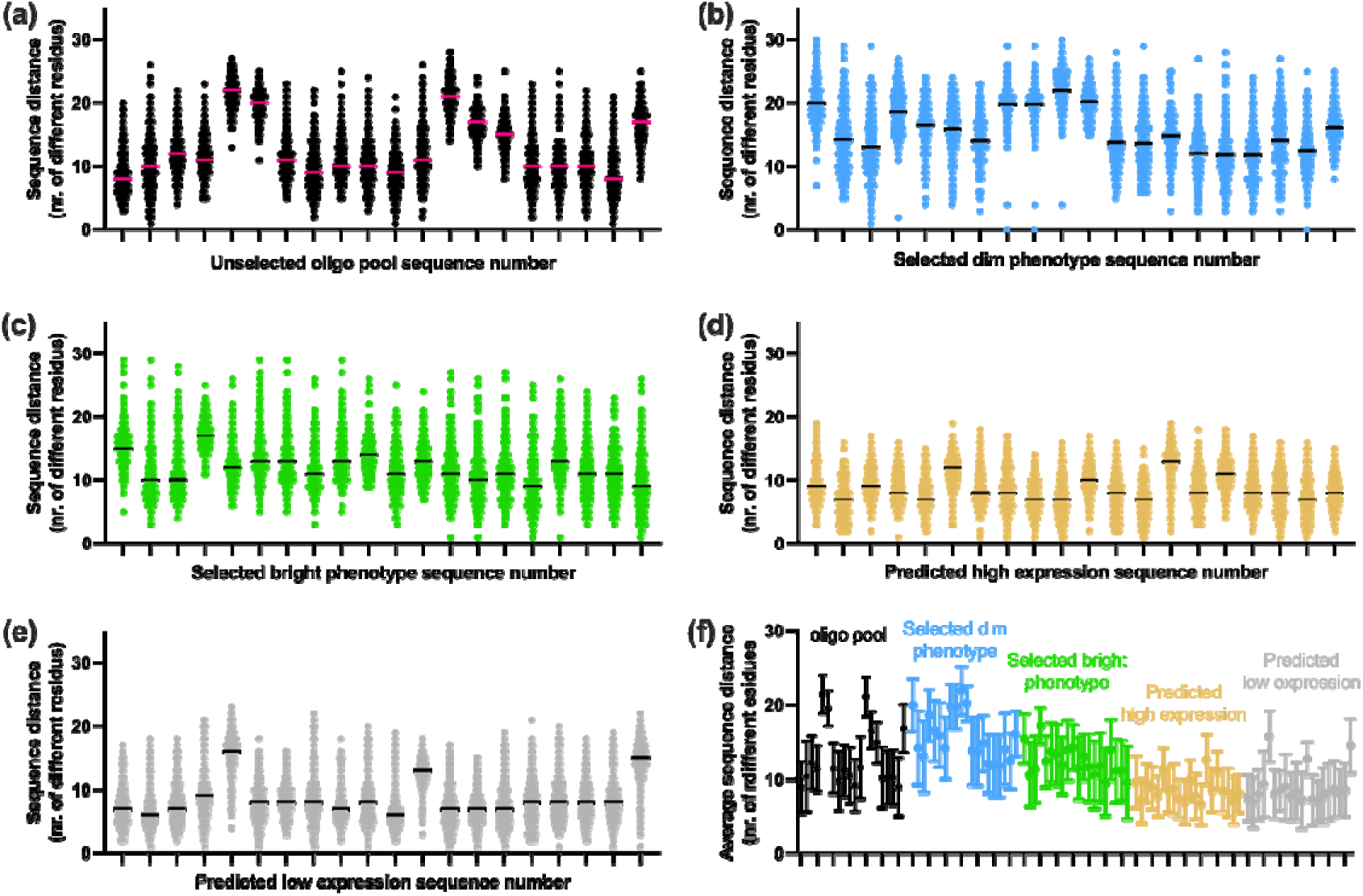
Between-sequence distances at the amino acid level for the different protein populations in this study. Two hundred sequences were selected arbitrarily from **(a)** unsorted and **(b, c)** FACS-sorted populations as well as from protein sequences predicted by ML to be high-or low-expressing **(d, e)**. Sequence distance was determined as percent ID for the first 20 proteins against the rest of the dataset after multiple sequence alignment with ClustalOmega, maintaining the input order in the output file. **(a)** The original design library has an average between-sequence distance of approx. 12 residues, although distributions are broad. **(b, c)** Selected library populations show similar sequence distances to the original library. **(d, e)** As expected, Inter-sequence distances are lower within a subset of proteins that share a strong prediction for their expression phenotype. **(f)** Data from panels **a-e** displayed on the same axes for direct comparison.

**Supplementary Figure S2.**
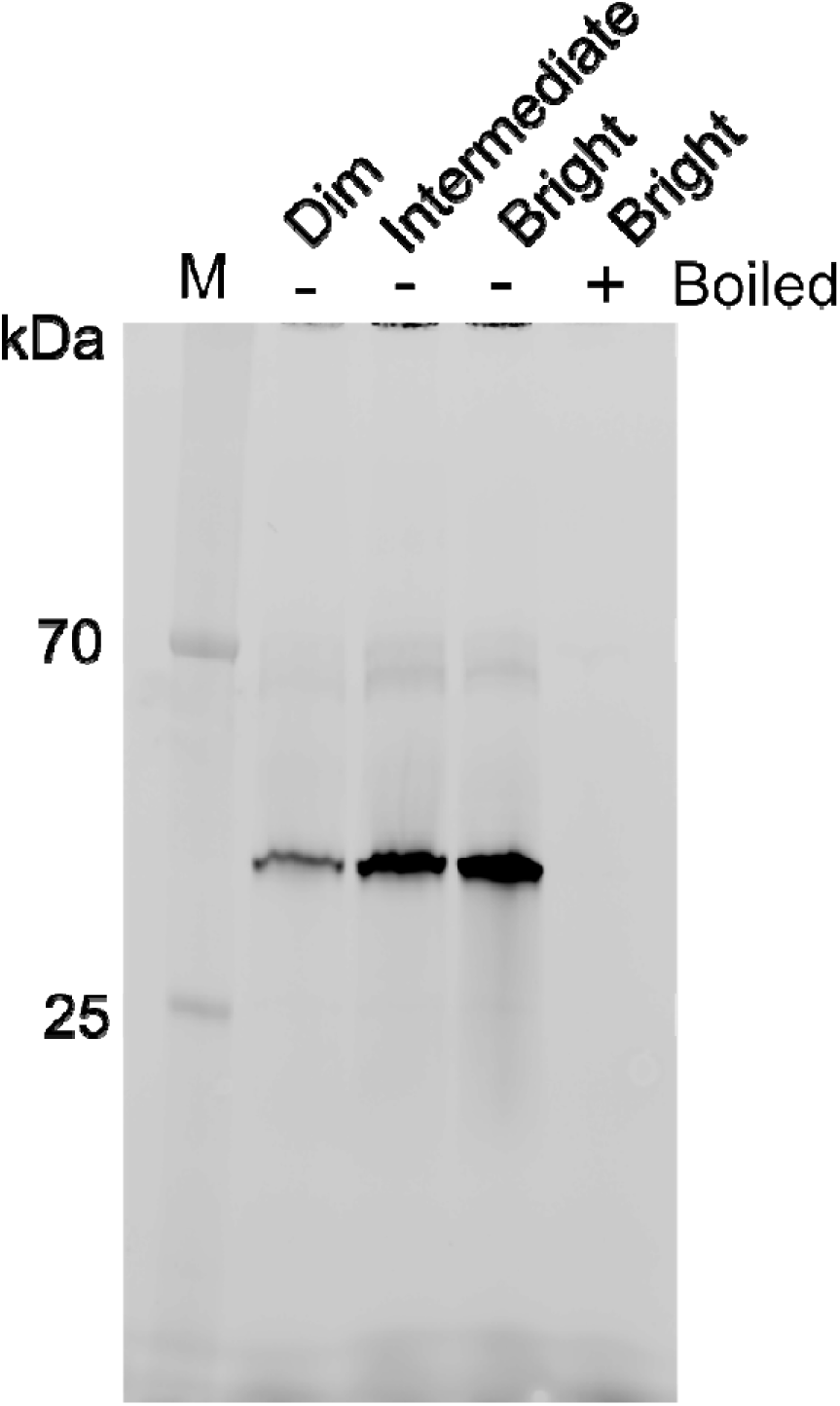
SDS-PAGE in-gel fluorescence of whole-cell lysates expressing individual proteins from the Dim and Bright sorted populations. Sample labelled ‘Intermediate’ is a protein from the Bright population with moderate bulk fluorescence level. Theoretical molecular weight of the fusion protein is 42 kDa.

**Supplementary Figure S3.**
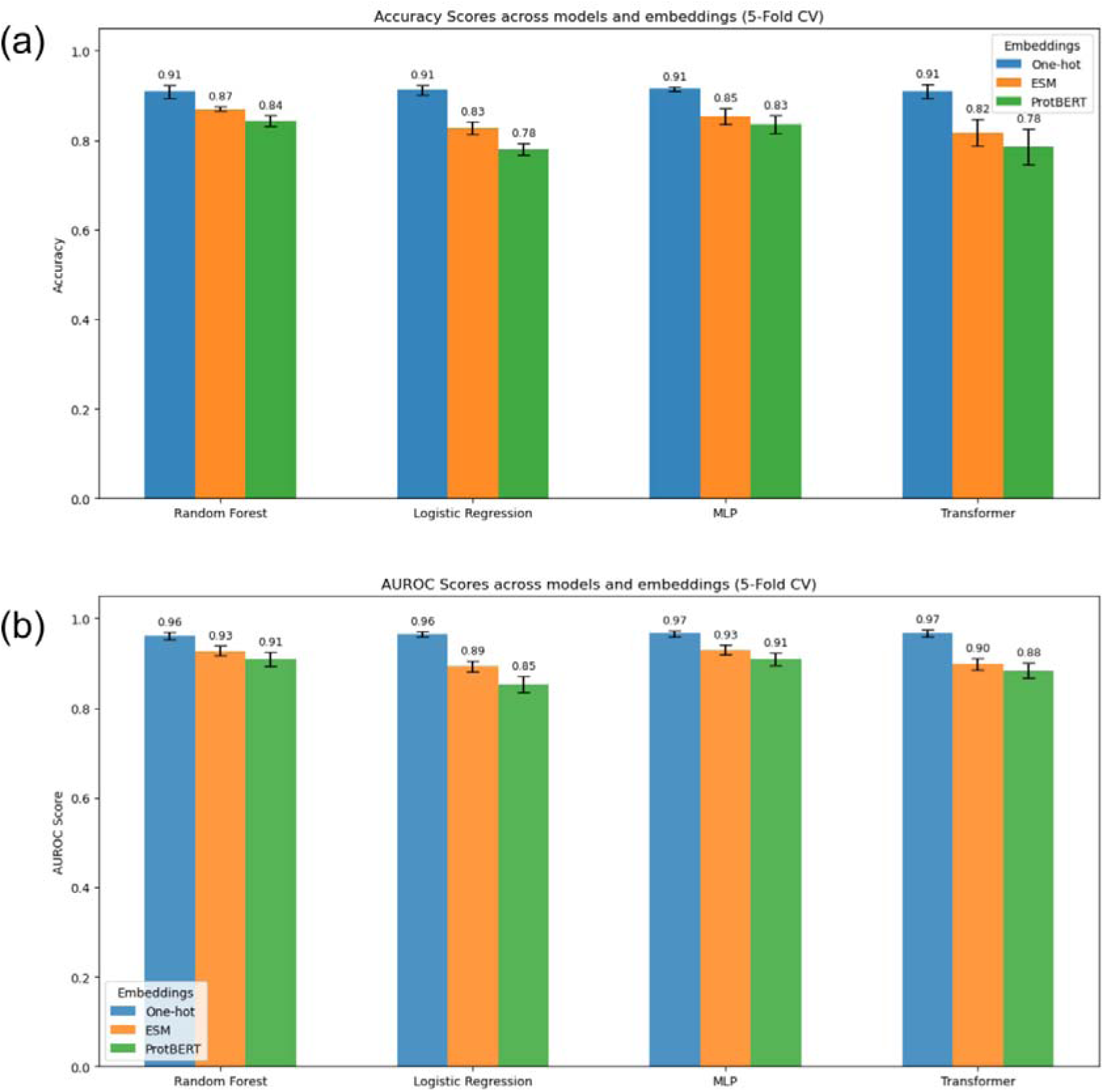
Comparison of four classification models trained on three amino acid sequence embeddings. **(s)** Accuracy score from each model. **(b**) Area under ROC curve (AUROC) Data are mean ± standard deviation in 5-fold cross-validation using the data specified in the main text.

**Supplementary Figure S4.**
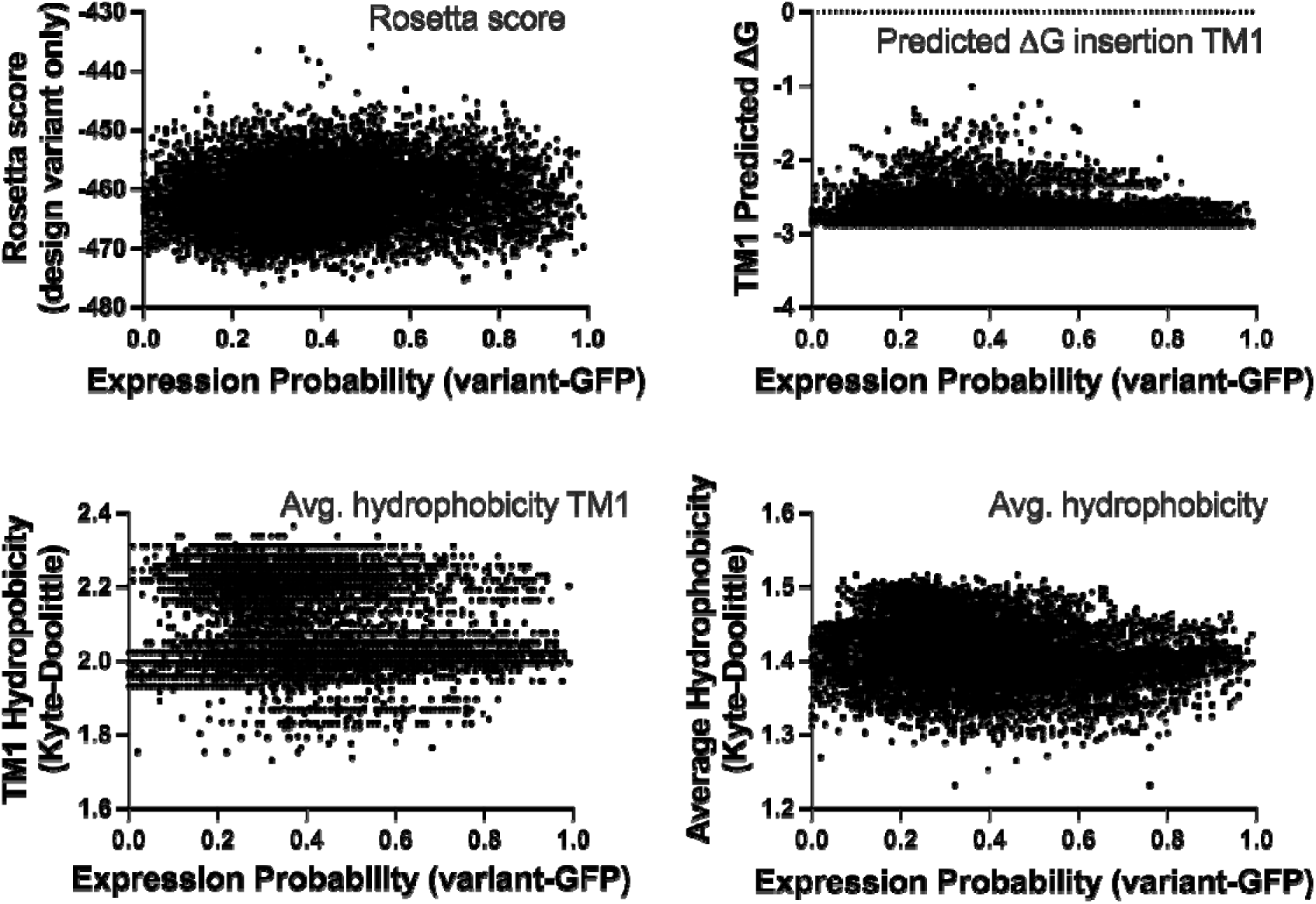
Simple biophysical sequence metrics do not correlate with expression probability from the machine learning model.

**Supplementary Figure S5.**
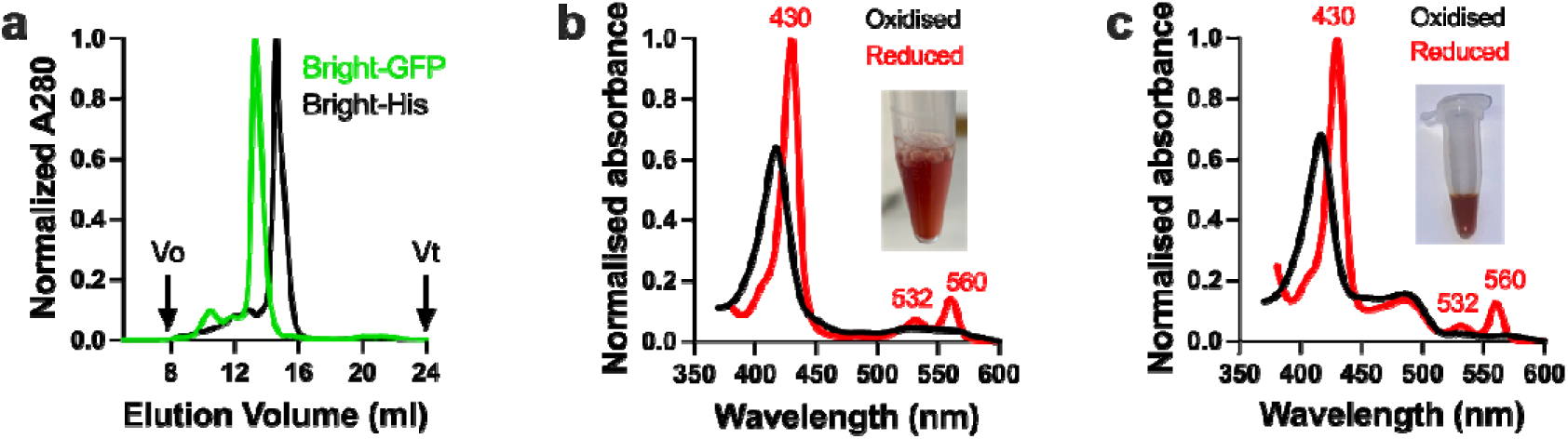
Studying the same ‘bright’ library variant with either GFP or His tag. (**a**) Size-exclusion analysis showing that the bright variant is monodisperse regardless of the purification tag used. Column is 10/300 S200 using 0.24% Cymal-5. (**b**) Absorbance spectroscopy of His-tagged protein, with photo of purified protein inset. (**c**) Absorbance spectroscopy of GFP-tagged protein, with photo of purified protein inset (same data as Fig. 3f in the main text).

**Supplementary Figure S6.**
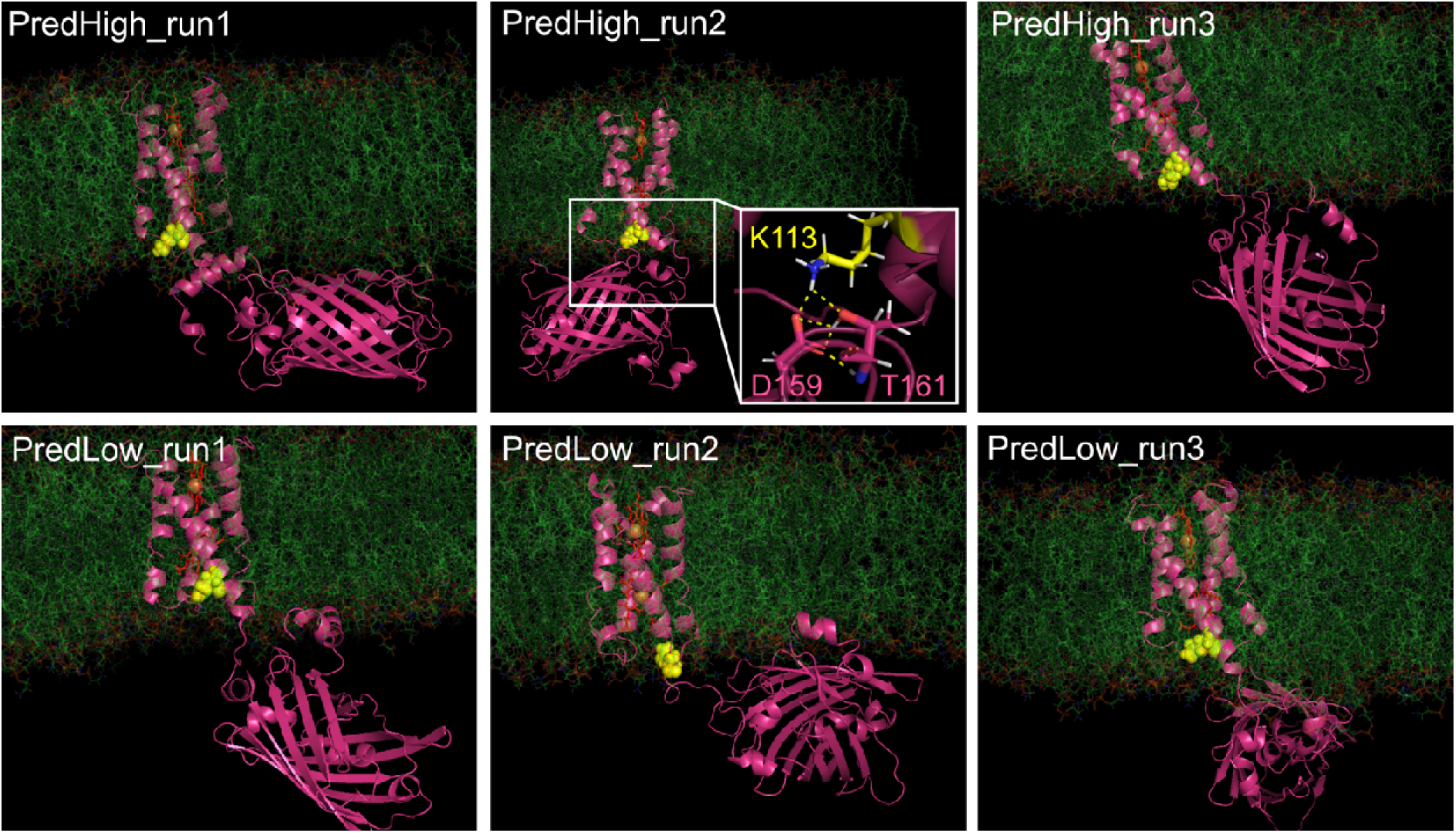
Molecular Dynamics simulations for two GFP-fused variants, PredHigh (K113) and PredLow (Q113). No clear interaction is observed between sequence position 113 (*yellow*) and the fused GFP, except for moderately persistent salt bridges formed in 25% of frames of one repeat (*PredHigh run2*). All data are the final frame of independent MD simulations over at least 300ns. Solvent hidden for presentation.

**Supplementary Figure S7.**
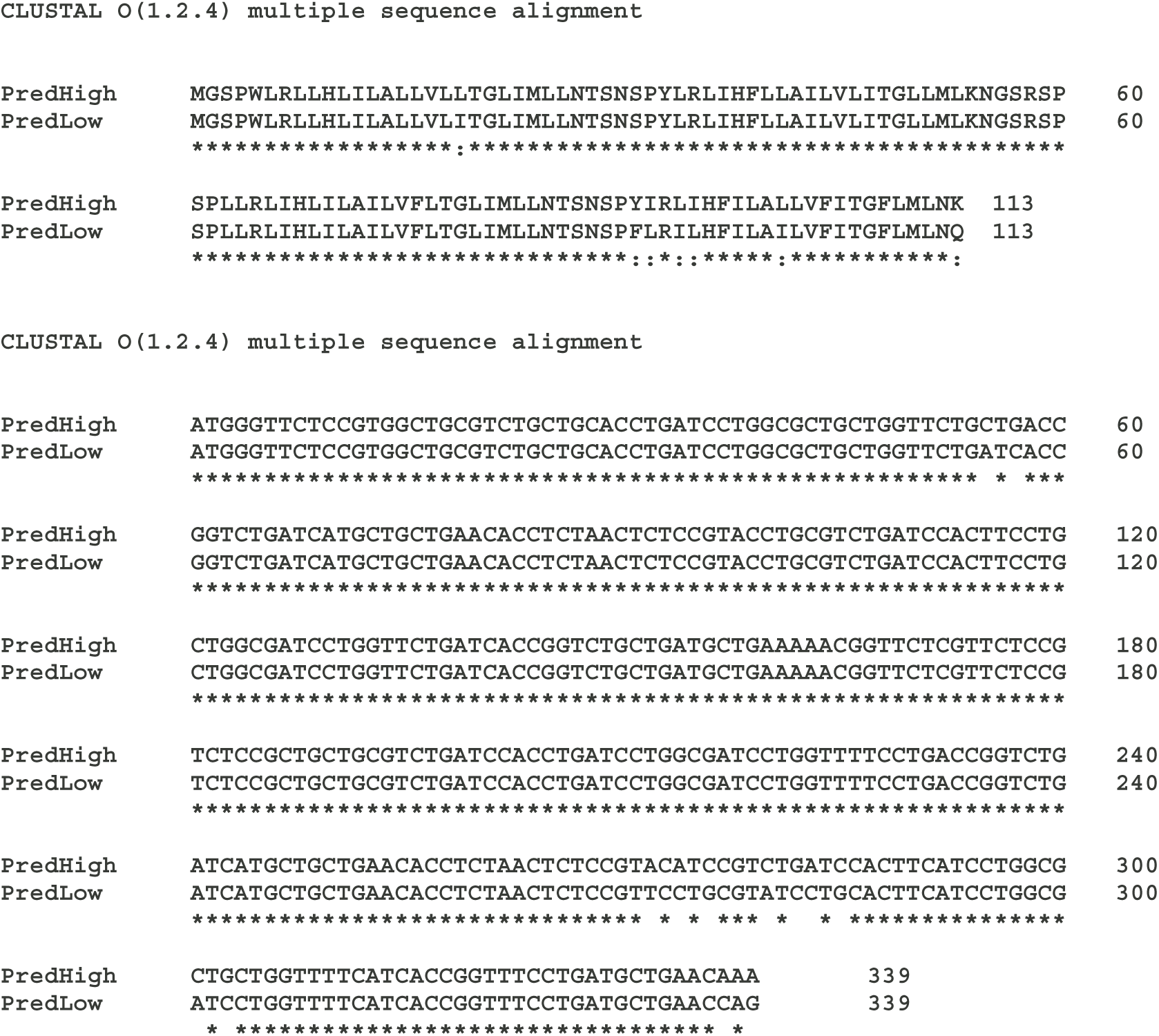
Multiple sequence alignment of two closely-related proteins with disparate expression outcomes. These proteins were used for expression engineering as shown in Figure 5.

**SupplementaryFigure S8:**
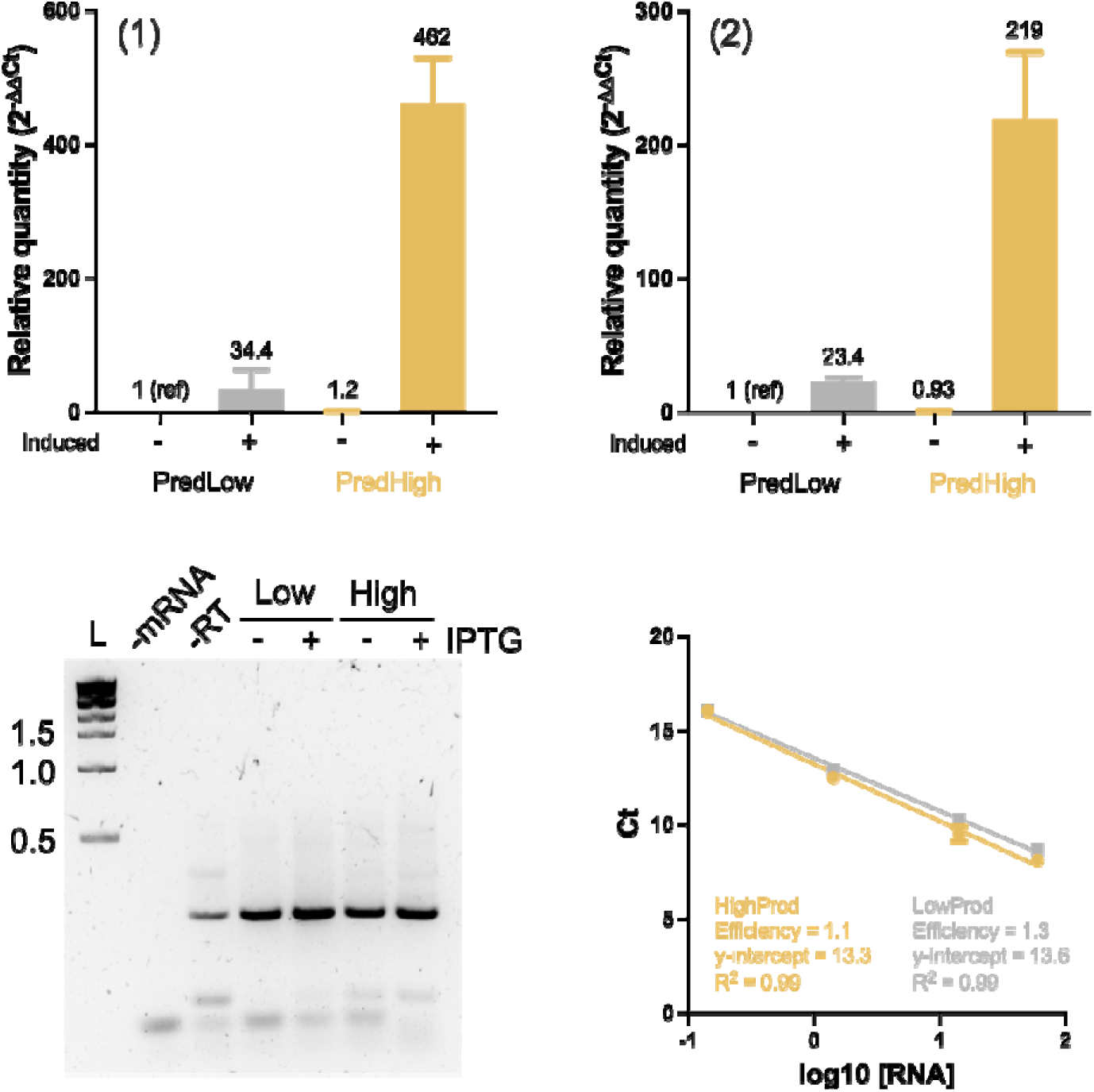
Results of RT-qPCR. Top panels show RT-qPCR results from two independent biological replicates; panel (1) is Fig. 5f in the main paper. In each case uninduced cells carrying variant PredLow are used as a reference. Post-reaction agarose gel shows an amplicon at the expected size of 239 bp, with minor contamination from DNA and primer products. RT-qPCR efficiency is ∼1.

## Notes

https://doi.org/10.6019/S-BSST2184

### Summary of Updates

Rewrote sections for clarity. Included new MP benchmarking data. Removed Figure 6 and associated speculative material to focus on the core findings of the work.

